# Whisker stimulation with different frequencies reveals non-uniform modulation of functional magnetic resonance imaging signal across sensory systems in awake rats

**DOI:** 10.1101/2024.11.13.623361

**Authors:** Jaakko Paasonen, Juha S. Valjakka, Raimo A. Salo, Ekaterina Paasonen, Heikki Tanila, Shalom Michaeli, Silvia Mangia, Olli Gröhn

## Abstract

Primary sensory systems are classically considered to be separate units, however there is current evidence that there are notable interactions between them. We examined the cross-sensory interplay by applying a quiet and motion-tolerant zero echo time functional magnetic resonance imaging (fMRI) technique to elucidate the evoked brain-wide responses to whisker pad stimulation in awake and anesthetized rats. Specifically, characterized the brain-wide responses in core and non-core regions to whisker pad stimulation by the varying stimulation-frequency, and determined whether isoflurane-medetomidine anesthesia, traditionally used in preclinical imaging, confounded investigations related to sensory integration. We demonstrated that unilateral whisker pad stimulation not only elicited robust activity along the whisker-mediated tactile system, but also in auditory, visual, high-order, and cerebellar regions, indicative of brain-wide cross-sensory and associative activity. By inspecting the response profiles to different stimulation frequencies and temporal signal characteristics, we observed that the non-core regions responded to stimulation in a very different way compared to the primary sensory system, likely reflecting different encoding modes between the primary sensory, cross-sensory, and integrative processing. Lastly, while the activity evoked in low-order sensory structures could be reliably detected under anesthesia, the activity in high-order processing and the complex differences between primary, cross-sensory, and associative systems were visible only in the awake state. We conclude that our study reveals novel aspects of the cross-sensory interplay of whisker-mediated tactile system, and importantly, that these would be difficult to observe in anesthetized rats.

## 1. Introduction

The perception by the senses of surrounding events and objects is one of the most important tasks undertaken by the brain^1^, as sensory information is the basis for decision-making, planning of future actions, and subsequent motor control^2^. The core structures of the different primary sensory circuits have now been well characterized with various electrophysiological, histological, and neuroimaging techniques. However, while most of the sensory research has focused on one sensory system at a time, there is persuasive evidence that sensory circuits form an integrated system at multiple levels of the ascending pathways^1,3–5^. For instance, the primary task of the whisker-mediated tactile system in rodents is to connect mechanoreceptors around the shafts of individual whiskers to the barrel region of the primary somatosensory cortex (S1bf) via nuclei in brainstem and thalamus^2,6^. The whisker-mediated tactile system, however, also has projections to various, likely multisensory regions, such as inferior colliculus (IC)^7^, superior colliculus (SC)^6,8,9^, medial geniculate nuclei (MGN)^8^, zona incerta (ZI)^6^, posterior parietal cortex (PPC)^10^, and secondary somatosensory cortex (S2)^6,11^. Due to the recent emphasis on cross- and multisensory perspective^3^ and findings, such that an auditory stimulus can induce or modulate neural activity in visual cortex or vice versa^3,12,13^, it does seem that many thalamic nuclei and parts of primary cortices should be considered as multisensory rather than specific to a certain sense^3,12,14,15^.

Due to their limited spatial coverage, it is challenging to undertake electrophysiological experiments to clarify the brain-wide cross-sensory interplay. Instead, modern neuroimaging techniques are ideal for undertaking whole-brain studies with relatively good temporal and spatial resolution. Indeed, several functional magnetic resonance imaging (fMRI) studies in human volunteers^16–25^ and non-human primates^26,27^ have explored the signal changes occurring in non-core sensory regions to unisensory stimuli, addressing such fundamental questions as which parts of the non-core sensory circuits are activated by an input into the core sensory circuit. However, more complex questions, such as how the non-core sensory circuits react to varying inputs into the core sensory circuit, have remained largely unexplored^28,29^. Moreover, most of the previous work has focused on the cortical interplay of auditory and visual systems, leaving other sensory systems and subcortical regions less extensively evaluated. Only a few fMRI studies in humans^20,23^, non-human primates^27^, and anesthetized mice^28^ have examined cross-sensory responses to tactile stimuli. Despite the whisker-mediated tactile system being one of the most commonly studied sensory systems, none of the previous whisker fMRI studies^30–40^ have adequately characterized cross-sensory activity in rats. Notably, the vast majority of the rat studies were done under anesthesia, which affects the organization of brain function^41^ and may thus hinder the detection of signal changes reflecting cross-sensory processing^28^. Lastly, the loud scanner noise inevitably present during traditional fMRI sequences represents a serious limitation in cross-sensory fMRI studies^17,42^, regardless of the target species.

Therefore, to understand better the mechanisms of sensory integration, we conducted controlled and quiet fMRI experiments in head-fixed awake and anesthetized rats to investigate the brain-wide activation patterns within and beyond the whisker-mediated tactile system. We hypothesized that if the neuroimaging approach was sensitive enough, a large amount of data should be able to reveal novel aspects of the cross-sensory interplay. Specifically, we aimed to 1) characterize the non-core brain-wide regions that respond to unisensory whisker pad stimulation, 2) study how the core and non-core regions respond to varying input into the whisker pathway, and 3) evaluate whether light anesthesia has a confounding effect on the detection of cross-sensory activity. To achieve this, we utilized a recently developed zero echo time (zero-TE) fMRI approach, Multi-Band SWeep Imaging with Fourier Transformation (MB-SWIFT)^43^, which is a quiet and movement-tolerant pulse sequence^44,45^, making it ideal for cross-sensory fMRI studies in awake animals. Furthermore, the distortion-free images^43,45,46^ and inherent 3-D radial acquisition strategy of MB-SWIFT provide high-quality images from cerebrum, regions near tissue-air interfaces, brain stem, and cerebellum^44^, making the zero-TE fMRI approach ideal for conducting brain-wide studies.

## 2. Results

To study the cross-sensory responses to unilateral whisker pad stimulation, 13 adult (7 males and 6 females) head-fixed Sprague-Dawley rats underwent 80 fMRI scans when they were either in the awake state or under light isoflurane-medetomidine (Iso+Med) combination anesthesia (Supplementary Table 1). Altogether, 1600 16-s stimuli blocks were analyzed (Supplementary Table 1). In order to allow a direct comparison between the awake and anesthetized states, a non-invasive mechanical air-puff mediated whisker deflection (typically 20-40°) was used in both conditions. Additionally, to explore further signal changes under a stronger stimulation approach, and to control the potential effect of the acoustic sound induced by the mechanical stimulation setup, electrical stimulation (2 mA biphasic pulse of 600 µs duration) via needle electrodes in the whisker pad was used under anesthesia. Throughout the manuscript, the term whisker pad stimulation is generalized to refer to either mechanical or electrical stimulation intending to evoke activity in the whisker-mediated tactile system.

To achieve varying inputs into the whisker-mediated tactile system, whisker pad stimuli were given at different frequencies (1, 5, 9, 13, or 17 Hz) in a randomized order. Natural exploratory whisking occurs over a wide frequency range (1-25 Hz), which can be roughly divided into slower large-amplitude (5-10 Hz)^6,47^ and faster small-amplitude (15-25 Hz)^6^ whisking. Our preliminary analysis indicated that fMRI responses to 5 and 9 Hz resembled each other, as did responses to 13 and 17 Hz stimuli (Supplementary Figure 1), which follows the categorization of different natural whisking types. Therefore, for simplicity in the analyses, the stimuli were categorized as either low- (1 Hz), mid- (5 or 9 Hz), or high-frequency (13 and 17 Hz) stimuli.

As there is marked variability in the terminology used in sensory research^48^, the terms used in this report are clarified here. The term “core” refers to the primary sensory system, i.e. the whisker-mediated tactile system. The term “non-core” refers to regions outside the whisker-mediated tactile system. The term “cross-sensory” refers to activity or interactions in non-core systems evoked by the input into the core system. The term “high-order” refers to cortical associative areas, while “low-order” refers to the other parts of sensory circuits up to the primary cortices.

### 2.1 Awake rats exhibit the largest whisker pad stimulation-induced signal changes outside the whisker-mediated tactile system

Group-level statistical maps (p<0.005, FWE-corrected) to mid-frequency whisker pad stimuli are shown in Figure 1, as this stimulation frequency range produced robust brain-wide activation pattern in all groups. Regardless of the stimulus approach or wakefulness state, signal changes were observed in regions covering the key nodes of the whisker-mediated tactile system, such as in ipsilateral principal and spinal trigeminal nuclei (Pr5/Sp5), in contralateral ventral posteromedial (VPM) and posterior (Po) thalamic nuclei, and in contralateral S1bf. A schematic illustration of the localization of these key nodes in the whisker pathway is shown in Figure 2A. Signal changes (Figure 1) were also detected in other parts of the whisker-mediated tactile system^6^, such as in S2, as well as in the lip region of primary somatosensory cortex (S1lip), likely indicating a non-whisker-related tactile sensory input from the snout during the stimuli.

**Figure 1.**
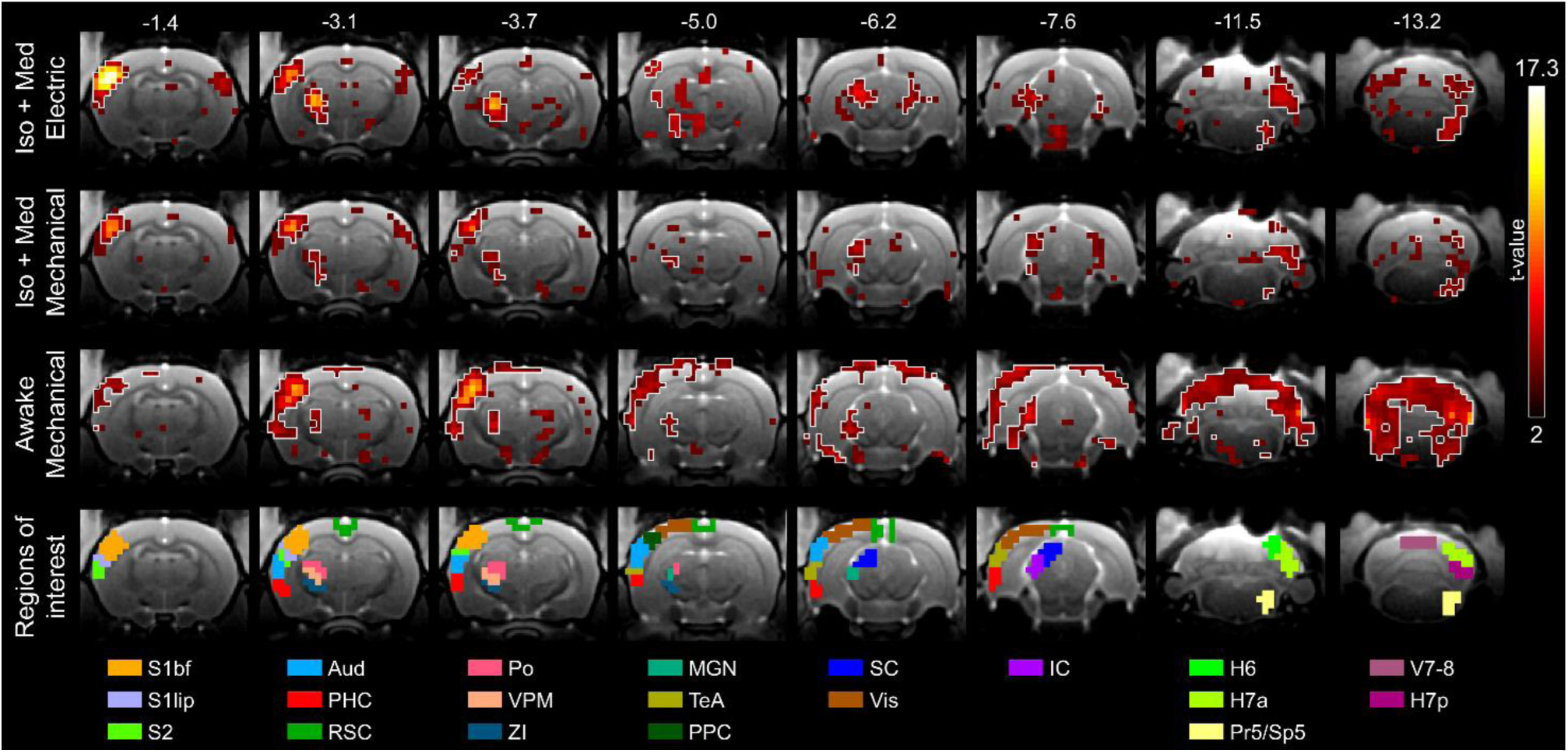
Group-level statistical maps to mid-frequency whisker pad stimuli. White outlines on the statistical maps indicate the regions with p < 0.005 (FWE-corrected), while the color maps are thresholded with t-value > 2. The results are obtained from 208-224 stimulus blocks during 26-28 scans in each group. Values in the top row indicate the approximate distance from bregma for each slice. Statistical maps are overlaid on high-resolution anatomical images. Aud, auditory cortex; H6, hemisphere of cerebellar lobule 6 (simplex); H7a, anterior hemisphere of cerebellar lobule 7 (crus 1 and 2); H7p, posterior hemisphere of cerebellar lobule 7 (paramedian 1); IC, inferior colliculus; Iso+Med, isoflurane and medetomidine anesthesia; MGN, medial geniculate nuclei; PHC, parahippocampal cortex;); Po, posterior thalamic nuclei; PPC, posterior parietal cortex; Pr5/Sp5, principal trigeminal nuclei and spinal trigeminal nuclei; RSC, retrosplenial cortex; S1bf, primary somatosensory cortex, barrel field; S1lip, primary somatosensory cortex, lip region; S2, secondary somatosensory cortex; SC, superior colliculus; TeA, temporal association cortex; V7-8, vermis of cerebellar lobules 7 and 8; Vis, visual cortex; VPM, ventral posteromedial thalamic nuclei; ZI, zona incerta.

**Figure 2.**
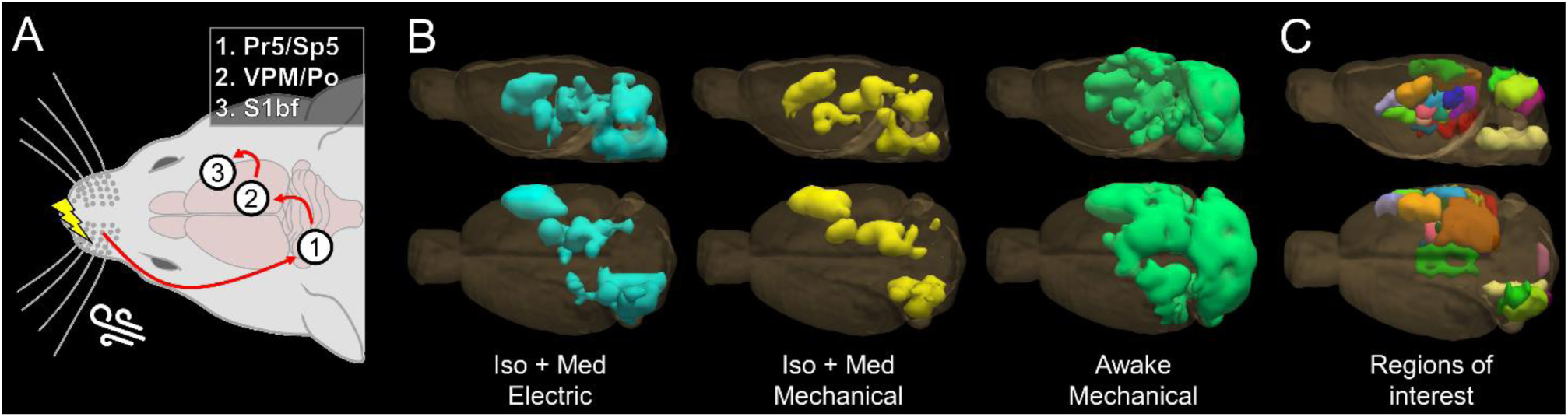
Schematic illustration of the whisker-to-barrel cortex core pathway activated by either mechanical (air puff) or electrical stimulus. **(A), a 3-D illustration of combined significant voxels (p<0.005, FWE-corrected) from low-, mid-, and high-frequency analyses within each group (B), and a 3-D presentation of regions-of-interest used in the subsequent analyses (C).** The results are obtained from 1600 16-s stimulation blocks given during 54 sessions (520-560 stimuli blocks per group). The colors for regions-of-interest in C correspond to those in Figure 1. Iso+Med, isoflurane and medetomidine anesthesia; Po, posterior thalamic nuclei; Pr5/Sp5, principal trigeminal nuclei and spinal trigeminal nuclei; S1bf, primary somatosensory cortex, barrel field; VPM, ventral posteromedial thalamic nuclei.

In addition to whisker-mediated tactile system, signal changes (Figure 1; p<0.005, FWE-corrected) were observed in regions covering the main nodes of the auditory pathways, such as in auditory cortex (Aud), MGN, and IC, and in visual areas, such as SC, in all groups. Furthermore, voxels covering ZI, which participates in the integration of sensory information and sensory-motor processes^49^, and many cerebellar regions, such as the hemisphere of lobule 6 i.e. simplex (H6), the anterior hemisphere of lobule 7 i.e. crus 1-2 (H7a), and the posterior hemisphere of lobule 7 i.e. paramedian 1 (H7p), displayed significant signal changes in all groups. However, signal changes near the PPC were observed only in awake rats whereas in anesthetized rats they were only evident after electrical stimulation. Moreover, in the awake rats, the statistical map partially covered the retrosplenial (RSC), visual (Vis), temporal association (TeA), and parahippocampal (PHC) cortices. Cerebellar signal changes were also clearly more widespread in awake rats, extending to additional regions, including the vermis of lobules 7 and 8 (V7-8) that has a representation of the face^50^, and generally indicating bilateral activity. In summary, awake animals expressed signal changes to tactile stimulation in large parts of the somatosensory, auditory, and visual systems up to cortex, in high-order regions, and also widely in the cerebellum. In anesthetized animals, the responses to stimuli were more restricted in comparison to those evident in awake animals, with notably absent signal changes in the high-order cortical regions and certain areas of the cerebellum.

3-D illustrations in Figure 2B and Supplementary Video 1 show the group-specific localization of significant signal changes (p<0.005, FWE-corrected) combined from low-, mid-, and high-frequency stimulation analyses. Altogether, signal changes were observed in 2234 unique voxels with mechanical stimulation in awake rats, in 729 unique voxels with electrical stimulation under anesthesia, and in 469 unique voxels with mechanical stimulation under anesthesia, further emphasizing the striking differences in spatial coverage of the signal changes between the awake and anesthetized conditions. Analogous 3-D illustrations of the statistical maps for each group and stimulation frequency range are shown in Supplementary Figure 2, suggesting that there are differences between awake and anesthetized rats in the regional fMRI responses to stimulation frequency-modulation.

To investigate further the frequency-dependency and temporal characteristics of fMRI responses in different brain areas, regions-of-interest (ROIs; Figure 1, Figure 2C, Supplementary Video 1) were derived based on 1) the localization of the activated areas (Figure 2B) in an anatomical atlas^51^ and 2) a literature search to reveal potential structural or close functional connections between the anatomical region and the whisker-mediated tactile system (see Introduction and Discussion). The list of selected ROIs and corresponding abbreviations are shown in Table 1, which also indicates that while all ROIs contained significant voxels (p<0.005, FWE-corrected) in awake animals, only 15/20 and 14/20 ROIs contained significant voxels with electrical and mechanical stimulation in anesthetized rats, respectively. The ROIs were first categorized into core and non-core regions, and further into five subgroups, namely whisker-mediated tactile system, high-order regions, auditory regions, visual regions, and cerebellum (Table 1).

**Table 1.**
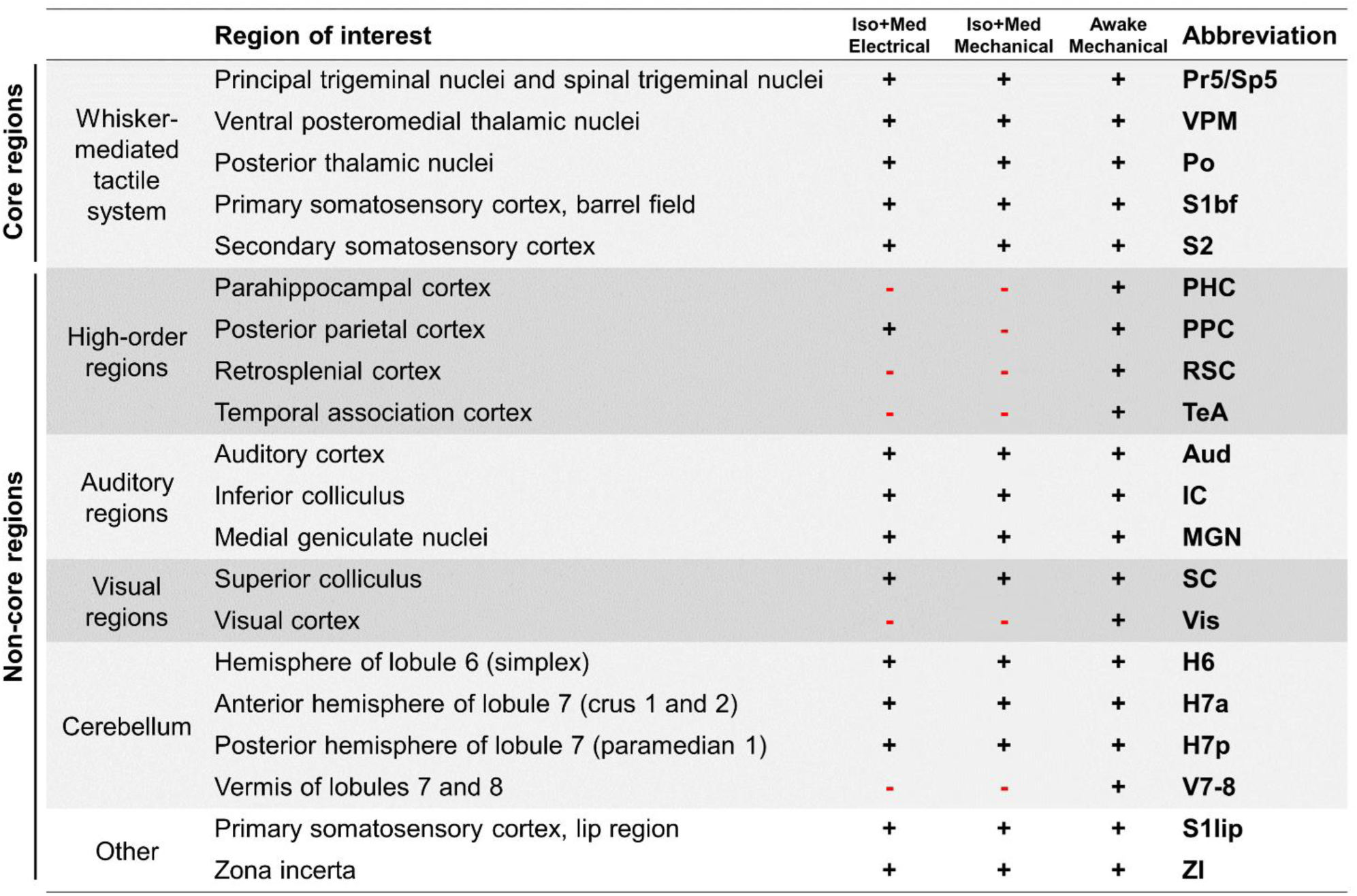
Regions of interest (ROIs) derived from the statistical maps and literature search. Awake animals exhibited signal changes (p<0.005, FWE-corrected) in all ROIs (indicated by +), while only 15/20 and 14/20 ROIs contained significant voxels in response to electrical or mechanical stimulation in anesthetized rats, respectively. In the subsequent evaluation, the ROIs were grouped into either core or non-core regions, and further divided into whisker-mediated tactile system, high-order regions, auditory regions, visual regions, and cerebellum. Iso+Med, isoflurane and medetomidine anesthesia.

### 2.2 The fMRI responses in the whisker-mediated tactile system increase with stimulation frequency in awake but not in anesthetized rats

To study the characteristics of fMRI signals during the whisker pad stimuli, group-level average time series (Figure 3) were derived from the selected ROIs (Table 1). Subsequently, the average fMRI responses were calculated, and significance and linearity for the slopes between the stimulation frequency and average fMRI response strength were estimated (Figure 4).

**Figure 3.**
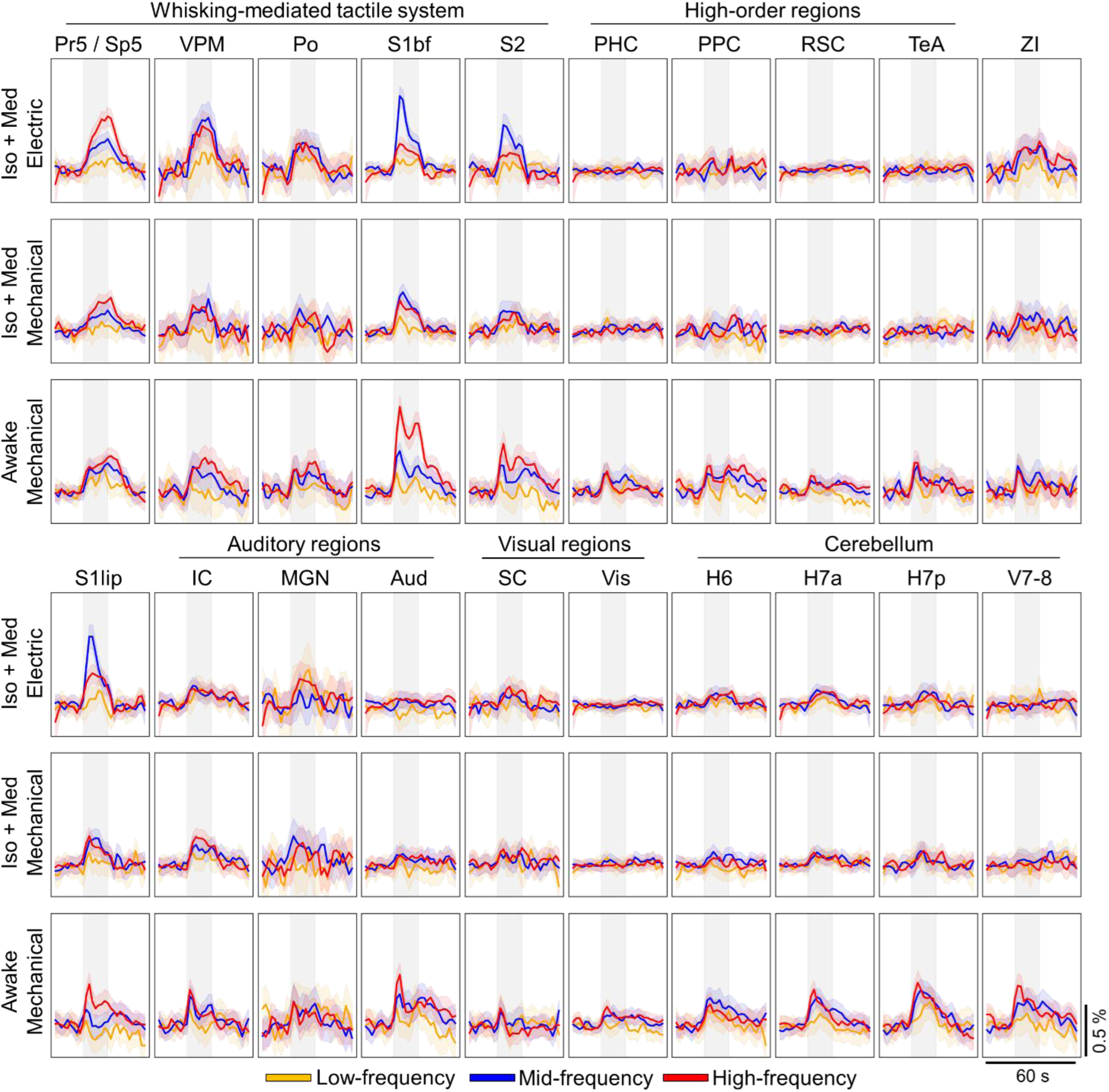
Group-level mean time series for each region-of-interest. Each average time series is a result of 104-224 stimuli (see Supplementary Table 1). The shaded vertical gray region indicates the timing for the 16-s stimulus block. The list of abbreviations for regions-of-interest can be found in Figure 1 and in Table 1. The 90% confidence interval is shown as a shaded region around the mean time series. Iso+Med, isoflurane and medetomidine anesthesia.

**Figure 4.**
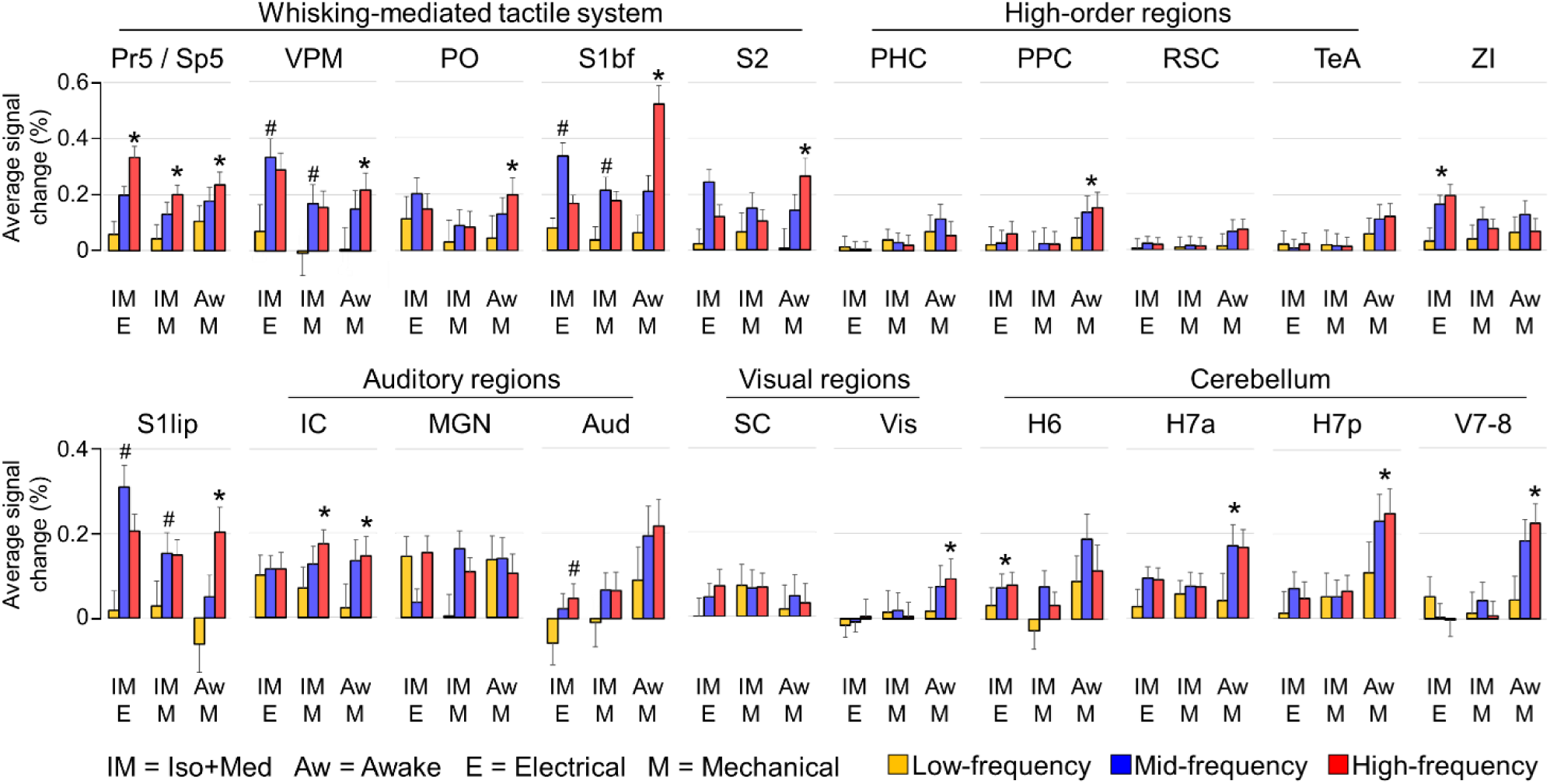
Group-level average fMRI response in each group and region-of-interest. The average response was typically calculated over a 20 s window starting from the stimulus onset (see Materials and Methods). For each region-of-interest and group, we tested 1) whether the slope determined by stimulation frequency and average response deviated from 0 (t-test), and 2) whether the relationship between the stimulation frequency and average response was linear (normality test for residuals of the fit). Asterisk (*) denotes a significant linear slope (t-test for slope p<0.05, normality test p>0.05), and hash (^#^) denotes a significant non-linear slope (t-test for slope p<0.05, normality test p<0.05). The full list for uncorrected p-values can be found in Supplementary Table 2. The error bars indicate the 90% confidence interval. The list of abbreviations for regions-of-interest can be found in Figure 1 and in Table 1. Iso+Med, isoflurane and medetomidine anesthesia.

When we examined the whisker-mediated tactile system, a significant linearly increasing response (t-test for slope p<0.05, normality test p>0.05) to low-, mid-, and high-frequency stimuli was detected in Pr5/Sp5 in all groups (Figures 3 and 4, Supplementary Table 2). This is a key observation, as it confirms that the input into the first relay station of the primary sensory circuit was linearly modulated with the applied stimulation protocol. Little to no adaptation should occur in trigeminal ganglion neurons^52^ and Sp5^39^ with stimulation frequencies up to 18-20 Hz (17 Hz highest in the current work).

Despite the brain stem showing linearly increasing fMRI responses in all groups, this was not the case in the thalamus (Figure 4, Supplementary Table 2), which is the next key node of the whisker-mediated tactile system (Figure 2A). While both thalamic nuclei, VPM and Po, responded linearly (t-test for slope p<0.05, normality test p>0.05) to the whisker pad stimulation frequencies in awake rats, the relationship between stimulation frequency and fMRI responses was non-linear (t-test for slope p<0.05, normality test p<0.05) in VPM in anesthetized rats (Figure 4). In Po, there was no significant slope between stimulation frequency and fMRI responses when the animals were anesthetized, revealing a confounding effect of anesthesia as also suggested earlier^2^. A similar but more pronounced pattern was seen in cortical somatosensory areas S1bf and S2, where the slope between stimulation frequency and fMRI response was linear in awake rats, but was either non-linear or non-significant in anesthetized rats. Interestingly, a significant and linear effect between fMRI responses and stimulation frequency was observed in ZI with electrical but not with mechanical stimulation (Figure 4), which might hint at nociceptive processing^53^ during electrical stimulation.

Taken together, the fMRI responses in all key nodes of whisker-mediated tactile system followed the stimulation frequency in awake rats. In anesthetized rats, both the thalamic and cortical regions showed saturated fMRI responses, peaking already with the mid-frequency stimulation. These observations highlight the confounding effect of Iso+Med anesthesia on the signaling in the whisker-mediated tactile system at the higher stimulation frequencies.

### 2.3 The fMRI responses in non-core regions are weak in amplitude

As depicted by the statistical maps in Figure 1, several regions outside the whisker-mediated tactile system showed highly reliable signal changes in response to the whisker pad stimuli. However, as seen in Figures 3 and 4, the signal changes in non-core regions were weak in amplitude. For example, the fMRI responses to mid- and high-frequency stimulation were typically 0.20-0.60 % in the key nodes of whisker-mediated tactile system (Pr5/Sp5, VPM, and S1bf) across the groups, while only 0.05-0.20% in the auditory, visual, and high-order regions. In the presence of anesthesia, the key nodes of whisker-mediated tactile system exhibited higher average response strengths in comparison to auditory (p < 0.001 with electrical and p = 0.039 with mechanical stimulation, two-sample Student’s t-test), visual (p < 0.001 with both electrical and mechanical stimulation, two-sample Student’s t-test), and cerebellar (p < 0.001 with both electrical and mechanical stimulation, two-sample Student’s t-test) regions. In awake rats, the average signal changes were smaller in the auditory (p = 0.016, two-sample Student’s t-test), visual (p < 0.001, two-sample Student’s t-test), and high-order regions (p < 0.001, two-sample Student’s t-test) but not in cerebellar regions (p = 0.081, two-sample Student’s t-test) as compared to the key nodes of the core pathway. These observations confirm one of our initial hypotheses that the signal changes in non-core regions in response to whisker pad stimuli are weak in amplitude but nonetheless can be reliably detected with larger data sets.

### 2.4 Responses in most non-core regions do not increase with higher stimulation frequencies

Another notable observation in Figures 3 and 4 regarding the non-core regions is that the majority of the auditory, visual, and high-order regions did not exhibit a significant slope between stimulation frequency and fMRI response strength. Among the high-order regions in awake rats, only PPC showed linear slope between the two variables, while in RSC, PHC, and TeA no significant slope was observed (Figure 4). In SC and MGN, which are typically considered as parts of the visual rather than the auditory pathways, there was no significant slope between the stimulation frequency and fMRI responses in any of the groups. A similar observation was made in Aud with mechanically stimulated awake and anesthetized rats. However, a significant linear slope was observed in IC with mechanical stimulation in both awake and anesthetized rats, but not with electrical stimulation. This may suggest that the response was triggered by the audible noise from the mechanical stimulation setup, possibly masking the cross-sensory observations in IC. Nevertheless, the results obtained with silent electrical stimulation indicate that the stimulus-induced responses in IC did not correlate with the stimulation frequency, similar to the observations in SC observed in all groups. In summary, the responses in several auditory and visual regions did not seem to follow the responses in the whisker-mediated tactile system, regardless of the state of wakefulness or type of stimulus. A similar observation was made in most of the high-order regions in awake rats.

In contrast to the auditory, visual, and high-order regions, most of the cerebellar regions in awake rats showed a significant linear slope between the stimulation frequency and the strength of the fMRI response (Figure 4). This differs from the results obtained in anesthetized animals, where the majority of the cerebellar regions displayed no relationship between input and response. These observations indicate that the interactions between cerebellum and whisker-mediated tactile system had been disturbed by the Iso+Med anesthesia.

### 2.5 The fMRI response profiles in awake rats within the whisker-mediated tactile system are coherent but differ from non-core regions

As it became apparent that the signal changes between core and non-core regions differed in many characteristics, we next examined more closely the similarities in the response profiles to different stimulation frequencies across the ROIs. For this purpose, a three-point response profile, namely the fMRI average response plotted against the stimulation frequency, was derived from the data shown in Figure 4. Subsequently, the response profiles were hierarchically clustered, with the results being shown in Figure 5. Only regions exhibiting significant signal changes (Table 1) were included in the clustering.

**Figure 5.**
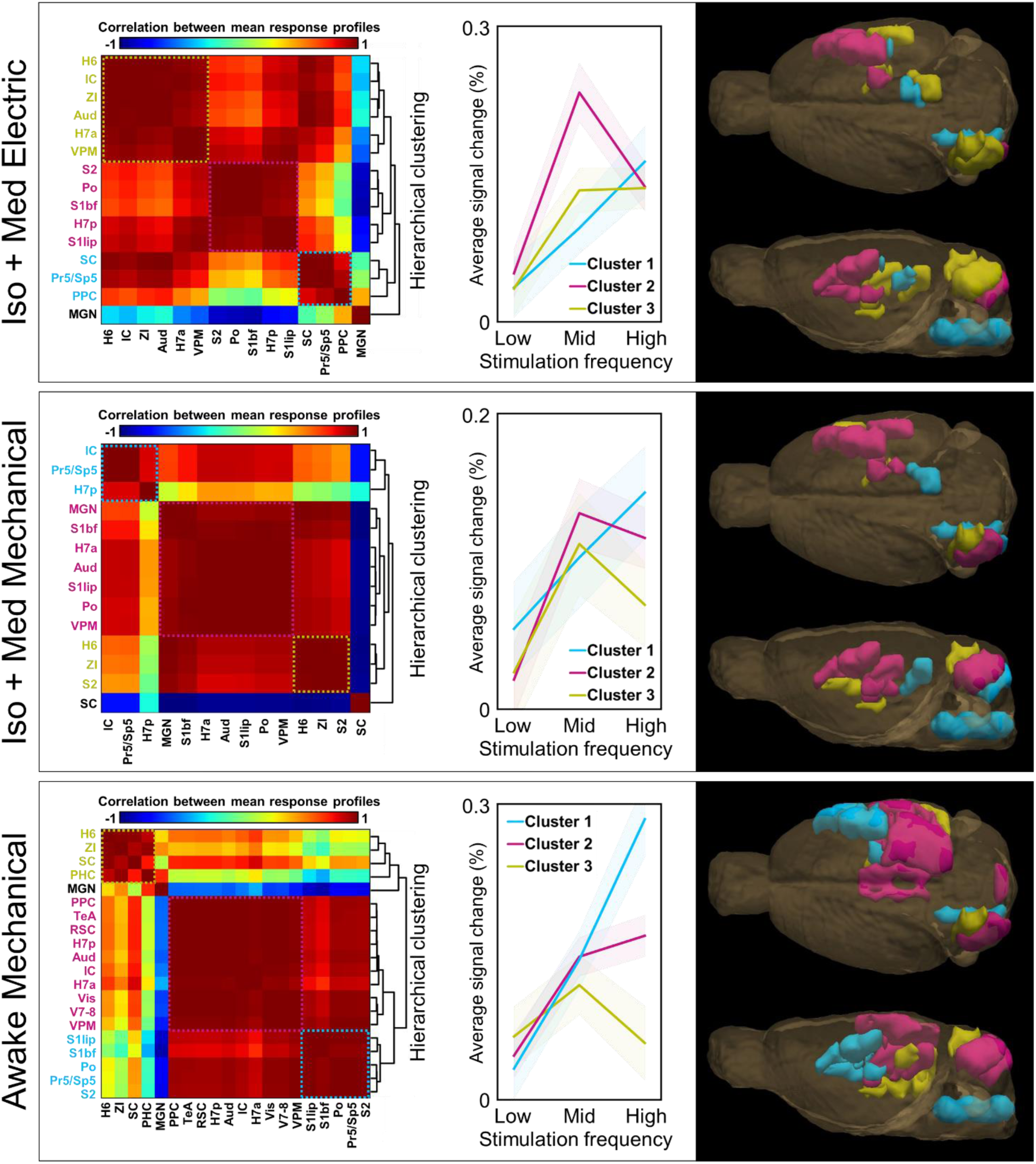
Hierarchical clusters of average fMRI response profiles. The average fMRI response profiles were derived from data shown in Figure 4, and clustered by using hierarchical clustering. The correlation matrices on the left indicate the similarity between the frequency response profile curves across regions-of-interest. The obtained hierarchical tree (or dendrogram) is shown on the right side of the matrices. Subsequently, average response curves were calculated for three main clusters for each group, which are shown in the middle. Clusters with single regions and high hierarchy were excluded from the illustrations, and regions without significant signal changes (Table 1) were left out from the analysis. The localization of the clusters is illustrated on the right. The clustered brain regions are color-coded in matrices, in the response profile graphs, and in the 3-D illustrations. The cluster including brain stem nuclei is color-coded as light blue in each group. The shaded region around the average response profiles indicates the 90 % confidence interval. The list of abbreviations for regions-of-interest can be found in Figure 1 and in Table 1. Iso+Med, isoflurane and medetomidine anesthesia.

In line with previous observations, clusters including Pr5/Sp5 (clusters #1, Figure 5) showed a linear increase in the response strength to the stimulation frequency in each group. In awake animals, the same cluster included most of the thalamic and cortical nodes of the whisker-mediated tactile system, while those were absent in the corresponding clusters in anesthetized animals. Despite the close spatial vicinity of many ROIs to the primary pathway in cortex and thalamus, there were no non-core regions in the same cluster with the primary pathway in awake animals. In awake rats, the non-core regions clustered separately with distinct frequency-modulation profiles in comparison to the primary pathway. Similar observations were not as evident in anesthetized animals, particularly after mechanical stimulation, where many thalamic and cortical nodes of the whisker-mediated tactile system were clustered together, but there were also many non-core regions included (clusters #2, Figure 4). The clustering results with electrical stimulation were more comparable with those obtained in awake animals, however, the response profiles were completely different.

These results strengthen the conclusion that the stimulation frequency-dependent modulation differs between core and non-core regions, possibly reflecting the fundamental differences between primary and cross-sensory sensory processing. It was difficult to draw the same conclusion in anesthetized animals, which suggests that Iso+Med anesthesia had exerted a confounding effect on signaling within and beyond the whisker-mediated tactile system.

### 2.6 The temporal dynamics of non-core regions differ from those in the whisker-mediated tactile system in awake rats

In addition to the spatial localization of the fMRI responses and the evaluation of the fMRI response amplitudes to different stimulation frequencies, we extended our analyses to study the temporal characteristics of the fMRI signals. An overview of the different types of temporal signal was obtained by undertaking a hierarchical clustering of the ROI-specific average time series (Figure 6), leaving out regions with no significant signal changes (Table 1). In anesthetized groups, the primary whisker pathway was split into two (mechanical stimulation) or three (electrical stimulation) clusters. Cluster #1, including the Pr5/Sp5, showed a slow rise and slow decay, while cluster #2, including S1bf, displayed a fast rise and fast decay, indicative of non-uniform temporal behavior in the primary pathway under anesthesia. In contrast, the correlation of temporal signal evolution was high between all key nodes of the whisker-mediated tactile system (Pr5/Sp5, VPM, and S1bf) in awake rats (cluster #1).

**Figure 6.**
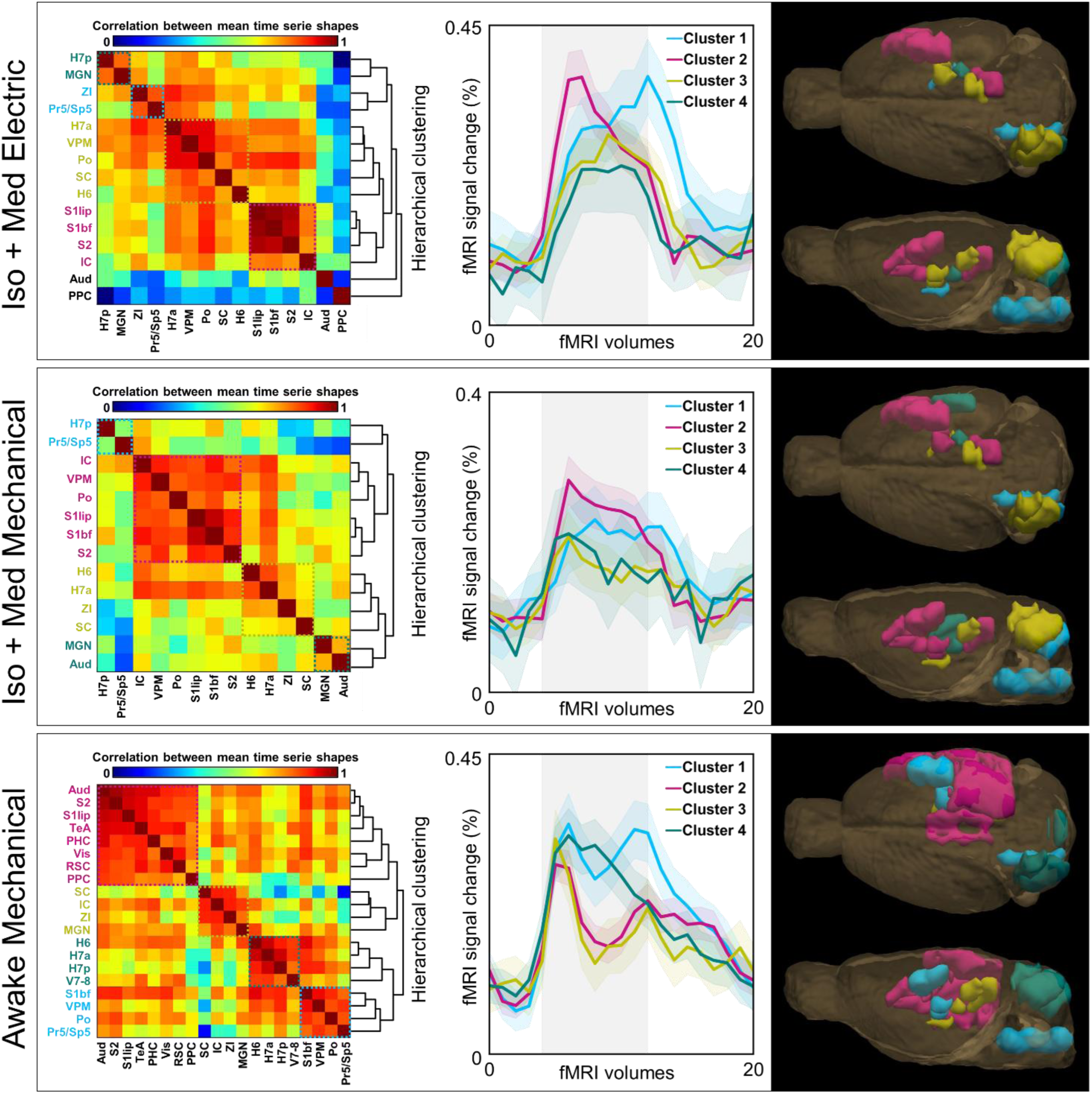
Hierarchical clusters of average fMRI time series. When studying the temporal characteristics of the signal changes, fMRI time series were averaged across all stimulation frequencies and clustered by using hierarchical clustering. The correlation matrices on the left indicate the similarity between time series across regions-of-interest. The obtained hierarchical tree (or dendrogram) is shown on the right side of the matrices. Subsequently, average time series were calculated for three to four main clusters for each group, which are shown in the middle. Clusters with single regions and high hierarchy were excluded from the illustrations, and regions without significant signal changes (Table 1) were omitted from the analysis. The localization of the clusters is illustrated on the right. The clustered brain regions are color-coded in matrices, in the response profile graphs, and in the 3-D illustrations. The cluster including brain stem nuclei is color-coded as light blue in each group. The shaded vertical gray region indicates the timing of the 16-s stimulus block. The shaded region around the average time series indicates the 90 % confidence interval. The list of abbreviations for regions-of-interest can be found in Figure 1 and in Table 1. Iso+Med, isoflurane and medetomidine anesthesia.

Overall, the categorization of ROIs into clusters based on their temporal profiles was vague in anesthetized animals (Figure 6). While with electrical stimulation cluster #2 represented mostly cortical, and clusters #3 and #4 thalamic and cerebellar temporal signals, there was no clear anatomical or functional basis to allow us to make a classification with mechanical stimulation. However, in awake rats, there was a clear separation for the primary pathway (cluster #1), cortical auditory, cortical visual, and high-order regions (cluster #2), subcortical auditory and visual regions (cluster #3), and cerebellum (cluster #4) based on the characteristics of their temporal signals. Interestingly, auditory, visual, and high-order regions exhibited a similar initial rise to core regions, but a fast decay to a lower amplitude level after only 4-6 s of stimulation, whereas a sustained high-amplitude response was maintained in the key nodes of whisker-mediated tactile system during the whole stimulus. Moreover, the cerebellar regions appeared to display a distinct profile compared to the regions in cerebrum and brain stem. These observations indicate that also the temporal dynamics of fMRI signal differ between the core and non-core regions during the sensory processing in awake rats. Importantly, these features would not be visible if the rats were subjected to Iso+Med anesthesia.

There were a few additional observations in the time series (Figure 6) worthy of consideration. First, the rise of the average signal to peak value appeared generally slower (4-8 s) under anesthesia in comparison to that in awake rats (2-4 s), which is in line with the earlier reports of a slower hemodynamic response function in the presence of anesthesia^54,55^. Second, the awake rats exhibited a biphasic fMRI signal with an initial peak and a delayed peak (except in cerebellum), which was not visible in anesthetized rats. Similar observations with different techniques have been reported earlier^56^. The later peak may originate from the slower components of functional hyperemia during prolonged stimulation^57,58^, which may be disturbed by the presence of anesthesia^59^. Third, the fMRI signal appeared to rise ∼2 s before the onset of the stimulus in awake rats, which may be a true response due to the learning and anticipation of the repetitive stimulus in awake animals^60^.

## 3. Discussion

As far as we are aware, this is the first study that has evaluated brain-wide cross-sensory activity to whisker pad stimulation in the rat brain, such as that detectable in the auditory and visual pathways. Moreover, it is also the first time that fMRI has been exploited to examine the stimulation frequency-independence of the responses in non-core regions. Importantly, we also demonstrated that even light anesthesia not only affected the evoked activity in the primary sensory pathway, but also disrupted the large-scale cross-sensory activity to a unisensory stimulus.

### 3.1 Brain-wide activity evoked by unisensory stimulus

The most typical and expected finding emerging from whisker pad stimulation fMRI studies has been the response in contralateral S1bf^30–39^. However, although a few other investigators have detected activity in other key nodes, such as in thalamus (VPM or Po)^38,39^ or brain stem (Sp5)^39^, none of the previous reports have described signal changes in cross-sensory regions, such as in the auditory and visual systems. A few factors could explain the different outcomes between the current and previous studies. First, in contrast to the present work, the echo planar imaging protocols used in previous experiments may not have been ideal for whole-brain fMRI because of the traditional acquisition of a limited number of coronal 2-D slices^61^ as well as issues with sensitivity to image distortions induced by magnetic field inhomogeneities. These limitations can restrict the spatial coverage to areas with a homogenous magnetic field, possibly leaving many of the regions detected here outside the field-of-view or with distortions. Second, previous work may have intentionally focused on a single sensory system or a single region. As a result, earlier data might have included signal changes in relevant brain areas that went unnoticed or unreported because of the absence of an initial hypothesis. Third, as observed in the present work, the signal changes in the non-core regions are weaker than those in core regions and may thus be difficult to detect with small sample sizes. Fourth, and partially related to the previous point, the conventionally used arbitrary statistical thresholding of neuroimaging results may lead to unintentional exclusion of interesting findings^62^, which can be particularly detrimental when evaluating the weaker cross-sensory signals.

Even though there are no previous rat fMRI studies reporting such widespread signal changes to whisker pad stimulation as described here, a recent experiment in mice^28^ shares many similarities with the current work. In addition to the cortical and thalamic key nodes of the whisker-mediated tactile system, Dinh et al. (2024) reported signal changes in the visual system (SC and Vis) and PPC to electrical whisker pad stimulation. Although not specifically reported or discussed by the authors, the activation maps also appear to cover parts of Aud^28^. These observations indicate that similar cross-sensory responses to whisker pad stimulation can be detected with fMRI in visual, auditory, and integrative systems in both mice and rats.

Several mechanisms have been suggested to explain the cross-sensory interplay. The potential structural connections allowing a cross-sensory information flow involve feedback projections from the high-order regions to the cross-sensory cortices, direct projections between the sensory cortices, an input from the thalamic sensory tracts to multiple primary sensory cortices, an input from the periphery to cross-sensory (or multisensory) thalamic nuclei, and corticothalamo-cortical tracts between different sensory cortices^13,17,18,63^. Even though we observed signal changes in relevant regions, we cannot distinguish the prevailing paths for cross-sensory information flow because of the scale of seconds in the temporal resolution and limitations related to the hemodynamic response function. Increasing interest in developing millisecond-scale and more direct imaging methods for neuronal activity may resolve these technical limitations in the future.

As the high-order regions, or association areas, are thought to be involved in the integration of sensory information and subsequent cognitive and motor functions^5,64^, the fMRI signal changes to whisker pad stimulation in these regions are not unexpected. The RSC is considered to be involved in the integration of multimodal sensory information and cognitive functions^65,66^. It has structural connections with the sensory cortices (Aud, S2, Vis), thalamus (MGN, Po), ZI, PPC, PHC, and brain stem^65–68^. PHC, including the perirhinal, postrhinal and entorhinal cortices, responds to multiple sensory stimuli^69,70^ and has structural connections with many of the sensory and associative areas listed above^66,70^. PPC has classically been considered as a high-order integration hub for modality-specific sensory information^4^, involved in wide range of cognitive, associative, and sensorimotor processes^10^. PPC is innervated by direct projections from the primary sensory cortices, including S1bf^10^. The exact physiological function of TeA, however, has not been characterized. TeA receives inputs from more than a hundred regions, including the several primary sensory regions^71^, which hints that it has an integrative role. Overall, the observed fMRI signal changes in physiologically relevant high-order regions indicate that a simple unisensory stimulus can elicit large-scale, detectable fMRI responses in associative networks throughout the rat brain.

The coordination of motor activity has been considered as a main task of the cerebellum, but there is emerging evidence for a significant role also in cognitive and affective functions^72^. Many cerebellar regions are specialized in sensory processing^73^ and can receive sensory input from multiple sensory modalities^74^. As mentioned earlier, the cerebellum is only rarely examined in rodent fMRI studies, and there are no fMRI reports of cerebellar responses to whisker pad stimulation in rats. Several regions in the cerebellar cortex, mainly in the lobules 6-7, receive an input from the whisker pad^50,73–76^, and electrophysiological activity in response to whisker pad stimulation has also been reported in these areas^73,76^. Therefore, the evoked cerebellar activity in H6, H7a, and H7p in the current study aligns well with the histological and electrophysiological findings. Interestingly, a recent study revealed bilateral projections from the trigeminal sensory nuclei complex to cerebellar cortex^74^, which might explain the bilateral fMRI responses observed in the current work. Cerebellum also receives inputs from the cerebrum, although the cerebro-cerebellar connections are poorly understood due to their multisynaptic organization^72^. Nevertheless, there is evidence that, e.g., V7-8 receive projections from the somatosensory and cingulate cortex^72^, which can explain the fMRI responses in V7-8 observed here. Another possibility is motor coordination^50^ occurring in head-fixed awake rats in response to stimulation.

Even though the signal increases were largely observed in whisker-mediated tactile system, it was somewhat surprising that we could not detect any activation in the primary motor cortex and dorsolateral striatum, which are parts of the sensorimotor system^6^. Although the head-fixed condition allows many simplified experimental designs^77^, it is clear that the whisker pad stimulation during a head-fixed condition does not correspond to natural whisking^78^, which may explain the missing activity. Furthermore, the electrical activity in different layers of the motor cortex during whisking is complex, evoking both increases and decreases in neuronal firing rates^9^. This can lead to a relatively small net effect in the total firing rate^9^ and only minor hemodynamic changes. Additionally, a recent study suggested that the neurovascular coupling in rat striatum would be more complex than previously thought^79^, which may also complicate the data interpretation in our case.

### 3.2 Stimulation frequency-dependent modulation and temporal characteristics of regional signals

Few fMRI studies have explored the signal changes to whisker pad stimulation in the primary sensory areas of rats with different stimulation frequencies. Studies done under anesthesia report increasing response amplitudes in S1bf up to 5 Hz (range of 1 - 10 Hz)^37^, 5 Hz (range of 1-20 Hz)^39^, or 12 Hz (range of 4-30 Hz)^32^ stimulation frequencies, after which a reduction in the strength of the fMRI response was observed. In awake rats, 40 Hz stimulation yielded stronger responses in comparison to those after 5 Hz^40^. These findings agree well with our results, as the highest response amplitudes in S1bf were observed after mid- and high-frequency stimulation (range of 1-17 Hz) in anesthetized and awake rats, respectively. Additionally, these findings indicate that anesthesia modulates the fMRI response profile to whisker pad stimulation, and the highest response amplitudes may not be related to the natural low-frequency whisker movement (4-12 Hz) as previously suggested^32,39^. Supporting evidence for this proposal has been obtained with laser Doppler flowmetry, where the maximal cortical blood flow to electrical whisker pad stimulation was achieved with 5 Hz under anesthesia, while a linear increase in blood flow was observed in awake rats up to a stimulation frequency of 40 Hz^80^. With respect to areas other than S1bf, Devonshire et al. (2012) reported saturated fMRI responses in VPM after 5 Hz, and linearly increasing fMRI responses in Sp5 up to 20 Hz under anesthesia, closely resembling our results.

Other interesting findings of the current work are related to the stimulation frequency-dependent modulation and temporal characteristics of fMRI signals outside the core pathway in awake rats. The fMRI responses in the core regions followed the whisker pad stimulation frequency almost linearly, while the majority of the auditory, visual, and high-order regions exhibited no or a non-linear dependence between stimulation frequency and fMRI response amplitude. We can speculate that the exact primary sensory information is thus not carried over to the cross-sensory regions, suggesting early and distinct processing mechanisms for the primary and cross-sensory information types. Whether these differ in terms of the specificity or encoding of the sensory information remains unclear and will require further electrophysiological studies. In addition, the stimulation frequency-modulation in the auditory, visual, and high-order regions resembled each other, which may indicate that the cross-sensory and associative processing share similar features that differ from the processing of the detailed primary sensory information.

As well as the distinct stimulation frequency-dependent response profiles, we observed different temporal signal shapes between core and non-core regions. The key nodes of whisker-mediated tactile system exhibited a sustained signal increase throughout the stimulation period, while the signal in non-core cerebral regions declined to a lower level shortly after the initial peak. This may indicate that adaptation to repetitive stimulus in cross-sensory and associative processing occurs more quickly than in the primary system, i.e. the cross-sensory processing may be more sensitive to changes rather than to the actual sensory input. The fMRI signal changes in the cross-sensory regions are likely explained by increasing neuronal firing rates, as these have been observed in the cross-sensory thalamic and cortical neurons^63^. However, the evoked cross-sensory activity has also been associated with the phase reset of β-γ band oscillations in non-core cortical networks^63^. A sudden, synchronized firing of a large neuron population may create a high, acute demands on energy metabolism, potentially manifesting a peak-like hemodynamic signal. Although the exact reasons for the changes in the hemodynamic signal in non-core regions remain unclear, the findings related to the temporal signal characteristics do seem to indicate that the processing strategies to unisensory stimulus are different in core and non-core regions.

As described above, cerebellum receives sensory input from multiple sources, and manages motor coordination and cognitive functions, leading to several layers of activity that may be difficult to interpret from hemodynamic signals. H7a, H7p, and V7-8 exhibited linearly increasing fMRI responses with elevations in the stimulation frequency, which may suggest that the underlying neuronal activity is at least partially driven by the sensory input from the primary sensory system. As H7a and H7p receive inputs from the whisker pad^73–75^ and V7-8 from the primary sensory cortices^72^, both pathways appear to be relevant. There were factors which differentiated the temporal signal profile in cerebellum from other regions, such as the slow and steady response decay before the end of the stimulus and the lack of a second peak (Figure 6, bottom row), which in the cerebral cortex presumably originated from delayed astrocyte-mediated hyperemia^57^. Despite showing some similarities in the stimulation frequency-induced modulation compared to the primary pathway, the early signal decay in the cerebellum may hint at the presence of faster adaptation mechanisms to repetitive stimulus. The absence of the second peak in the response shape may be due to several factors, such as a lack of sustained neuronal activity, different processing strategies, or distinct mechanisms of functional hyperemia between the cerebellum and cerebrum.

### 3.3 The impact of anesthesia on cross-sensory and high-order processing

To date, no fMRI study has characterized or compared cross-sensory brain-wide activity in the awake and anesthetized conditions. fMRI responses to visual stimulation have been reported in cortical auditory areas in both alert and anesthetized monkeys, being larger in the alert condition^26^. Similarly, we observed cross-sensory signal changes to whisker pad stimulation in Aud in both anesthetized and awake rats, with the responses being stronger in awake animals. But in contrast to previous reports, our whole-brain approach allowed us to conduct a brain-wide evaluation of the effects of anesthesia on sensory responses.

Overall, the results obtained in awake and anesthetized animals exhibited many similarities. Under both conditions, whisker pad stimulation elicited responses in the key nodes of the whisker-mediated tactile system, in certain parts of auditory and visual tracts, and in cerebellar regions receiving inputs from the whisker pad. These regions may thus comprise the core structure for the low-level sensory processing of whisker-mediated tactile information, where cognition, attention, and decision-making play a minimal role. It is worth noting that the signal changes in Pr5/Sp5 were very similar under both conditions, suggesting that sensory input is perceived at the lowest level with little to no influence from the Iso+Med anesthesia. In line with the previous work, our data support the concept that certain aspects of cross-sensory processing can be reliably detected under anesthesia.

Nevertheless, there were multiple observations indicating that anesthesia can seriously mask brain-wide responses to sensory stimulation. First, in the anesthetized animals, many regions, particularly among the high-order regions, showed no signs of evoked activity. Second, similarly to previous observations^26^, anesthesia appeared to restrict fMRI responses to a smaller area compared to measurements in conscious rats. Third, anesthesia distorted the linear stimulation frequency-dependent behavior of fMRI responses along the primary sensory tract, exhibiting non-linear relationships that were not present in the awake state. Fourth, because of the disrupted linearity of the fMRI responses to the stimulation frequency under anesthesia, the average response profiles across regions lacked details, making it impossible achieve a physiologically relevant clusterization. And fifth, this was also the case when we examined the temporal signal characteristics; the measurements conducted under anesthesia lacked details preventing a physiologically relevant clusterization. The last three effects are perhaps the most crucial ones, as based on the anesthesia data alone, it would not have been possible to derive the main conclusion that the core and non-core regions express distinct stimulation frequency-induced modulation and temporal characteristics of fMRI signals.

As anesthesia is considered to disrupt integrative processes^81^, the lack of fMRI responses in the high-order regions (RSC, PHC, and TeA) in anesthetized rats is not surprising. If the cross-sensory information flow is dependent on the feedback projection from integrative regions or cortico-cortical projections, it may also explain the smaller evoked cross-sensory activity in Aud and Vis in the presence of anesthesia. However, PPC responded to electrical whisker pad stimulation in the anesthetized rats, as has also been reported in mice^28^, which indicates that certain associative regions may show a response to during strong stimulation even in the presence of anesthesia. Interestingly, V7-8 was the only cerebellar region showing no significant signal changes to stimulation in the anesthetized animals. As mentioned earlier, it is likely that V7-8 receives whisker-mediated sensory inputs from the cerebral cortex^72^, which indicates that cortico-cerebellar circuits, or prior integrative processes, are suppressed by anesthesia. Alternatively, the signal changes observed in V7-8 may be related to a coordination of muscle activity^50^ during the awake state, in response to the sensory stimulus. Despite being weak in amplitude, the responses in H6, H7a, and H7p appeared robust under anesthesia, which is also supported by the fact that neurons in H7a were reported previously to respond to electrical whisker pad stimulation in both awake and anesthetized mice^73^.

### 3.4 Limitations

Neurovascular coupling may vary along the whisker-mediated tactile pathway^39^. Nevertheless, we assume that the region-specific stimulation frequency-modulated change in the fMRI signal represents a qualitative change of the underlying neuronal activity. Moreover, assuming a roughly similar neurovascular coupling within a given larger region, we can functionally differentiate, e.g., two cortical structures based on their signal behavior. Comparisons for signal change amplitudes were done mainly to evaluate the detectability of the signal.

The mechanical air-puff stimulation introduced a sound source overlapping with the stimulus paradigm. The potential effect on the findings was minimized by using ear plugs in both anesthetized and awake rats, and as low air pressure as practically feasible to achieve whisker deflections. Furthermore, experiments with mechanical stimulation were partially controlled with the implementation of inaudible electrical stimulation. Indeed, we observed activation of the auditory tract in response to both electrical and mechanical stimuli. It has been reported that awake mice exhibited bilateral auditory responses to a noisy unilateral mechanical whisker pad stimulation apparatus^82^, whereas only unilateral auditory activity, which corresponds better to specific cross-sensory activity, was observed in the current study. Nevertheless, we cannot fully exclude the potential confounding effect of audible noise on the data obtained with mechanical stimulation.

Another limitation is the fact that rats were repeatedly anesthetized with isoflurane. As single surgical-level isoflurane anesthesia exposure can affect the brain for weeks^83^, it is possible that the anesthesia may have induced some unknown long-term effects on cerebral function.

In addition to the certain level of inherent spatial and temporal inaccuracy of hemodynamic responses, the isotropic spatial resolution of 625 µm in our functional images poses certain limitations. Because of the resolution, the analyses were restricted to larger anatomical areas, leaving out potentially relevant areas at the scale of a few voxels. Additionally, only small parts of the large anatomical regions may be participating in the multimodal activity^3^, which subsequently weakens our ROI-based signals when large ROIs are used. However, there is no detailed information available related to the exact intra-regional multimodal zones in the rat brain, which would help to focus these kinds of analyses.

Lastly, a few factors may explain why the fMRI signals in the current work appear weak in amplitude. First, the are differences in the origin of functional contrast between blood oxygenation level dependent measurements and zero-TE fMRI methods^43^, leading to different absolute response amplitudes but similar functional contrast-to-noise ratios^44^. Second, it has been suggested that whisker pad stimulation may be one of the weaker physiological stimuli used in preclinical fMRI, leading to signal changes as low as 0.3% even with the conventional fMRI techniques^34^.

## 4. Conclusions

Where applicable, our results are in good agreement with previous work. As novel findings, we demonstrated that whisker pad stimulation elicited weak but robust brain-wide cross-sensory activity in rats. Additionally, the non-core systems responded to stimulation in a very different way than the primary sensory system, likely reflecting the different encoding and processing modes for the primary, cross-sensory, and high-order information. Lastly, while low-order sensory activity could be reliably detected under anesthesia, high-order processing and complex differences between sensory systems were visible only in the awake state. Further studies will be needed to explore whether similar or different results can be obtained with different species, other types of stimuli, multisensory stimuli, varying stimulation intensities, and more complex stimulation protocols.

## 5. Materials and methods

### 5.1 Animals

Animal procedures were approved by the National Animal Experiment Board and conducted in accordance with the European Commission Directive 2010/63/EU guidelines. A total of 13 adult (250-430 g at the onset of the first fMRI scan) Sprague-Dawley rats (Envigo RMS B.V., Horst, Netherlands; seven males and six females) were used in 54 fMRI sessions to obtain data during 1600 stimuli blocks. No data needed to be excluded from the analyses. On average, one rat underwent four fMRI sessions, two of which were in awake state and two under anesthesia. Details related to weight, age, and number of experiments under different conditions are listed in Supplementary Table 1.

Rats were group-housed prior to the procedures, but individually housed during the experiments to minimize damage to the head-implant and the surgical wound. Rats were maintained on a 12/12 h light-dark cycle at 22 ± 2 °C with 50% − 60% humidity. Food and water were available *ad libitum*. All experiments were carried out between 8 a.m. and 5 p.m.

### 5.2 Head-fixation implant surgery

In order to obtain high-quality fMRI data during awake imaging without significant movement artefacts, a chronic implant for head-fixation was attached on the top of the skull of the rats. The implant was manually shaped to be compatible with a single-loop 22-mm inner diameter MRI coil. The surgical procedures were similar to that described earlier^44,45^. Briefly, the rat was anesthetized with isoflurane (Attane vet 1000 mg/g, Piramal Healthcare UK Limited, Northumberland, UK; 5% induction and 2% maintenance) in 30/70% O_2_/N_2_ carrier gas. The scalp was removed from the top of the skull, and anchoring screws were inserted on top of the olfactory bulb and cerebellum. Four tungsten wire electrodes were placed bilaterally over S1bf for electrophysiological recordings, but data from these were not included in the current study. Subsequently, layers of bone (Palacos R + G, Heraeus Medical, Hanau, Germany) and dental (Selectraplus, DeguDent GmbH, Hanau, Germany) cements were applied while leaving two pins (2-mm diameter) covered with heat-shrinkable tubes passing through the dental cement. After the hardening of the cement, the pins were removed, leaving two holes in the implant for head-fixation points. Buprenorphine (0.03 mg/kg s.c.; Temgesic, Indivior Europe Ltd, Dublin, Ireland), carprofen (5 mg/kg s.c.; Rimadyl, Zoetis Belgium SA, Louvain-la-Neuve, Belgium) and saline (10 ml/kg/day s.c.) were given twice a day for two days for analgesia and rehydration. After the surgery, rats were allowed to recover for 1-3 weeks before starting the habituation protocol.

### 5.3 Habituation protocol for awake measurements

The 2-week habituation protocol shared many similarities with our previously described protocols^44,45,84^. Briefly, 1-3 rats per day were gradually habituated to body restraint and head-fixation. During the beginning of the first week, the rats were allowed to familiarize themselves with the handling person, the sounds of the fMRI sequence, short (≤ 60 s) head and body immobilization times, and positive rewards (almond flakes or 1% sucrose water). During the 5^th^ day of the first habituation week, the rat was gently immobilized within an elastic plastic foam and a custom-made head-fixation apparatus and kept in the imaging holder for 10 min^84^. Light isoflurane maintenance anesthesia (1.3-1.8%) was used while inserting ear plugs and covering in the plastic foam, and while positioning or removing the rat from the imaging holder. After a pause for the weekend, the habituation was continued by gradually increasing the habituation time in the imaging holder from 10 to 25 min. Additionally, the rat was familiarized with the masking tape on the whiskers and the air puffs used to induce the mechanical whisker movement.

### 5.4 Whisker pad stimulation protocols

For the stimulation of whisker-mediated tactile system, either electrical stimulation of the whisker pad or an air puff-based mechanical deflection of the whiskers was used. Stimuli were given with a stimulus generator (STG4008-16mA, Multi Channel Systems MCS GmbH, Reutlingen, Germany; MC_Stimulus II software V 3.5.11), and performed unilaterally on the left or right with the general aim to produce equal amounts of data for both sides.

Electrical stimulation was used only in anesthetized rats to avoid the potential discomfort induced by the needles and stimulation current during awake state. Stainless steel 30G needle electrodes were placed between whisker row pairs A-B and C-D. A single electrical stimulation pulse consisted of 2 mA for 300 µs followed by −2 mA for 300 µs.

In the air puff-based mechanical stimulation, posterior-to-anterior deflections were achieved with 5-ms puffs of air directed at a piece of adhesive tape on the whiskers. The tape on the whiskers was approximately 20x15 mm in size and attached to a maximal tuft of the whiskers. The air puffs were produced with pressurized air regulated to 1.0-1.5 bar and gated by a solenoid valve (custom-made by Neos Biotec, Pamplona, Spain) that opened based on the signals received from the stimulus generator. The air from the solenoid valve was directed to the posterior side of the whiskers via a 7-m pneumatic pipe (6-mm inner diameter) and a 50-cm flexible tube (3-mm inner diameter). To achieve a reliable deflection of the whiskers (typically 20-40°), the angle of the tube end was fine-tuned for each experiment separately. Slow-motion video recordings in bench tests confirmed that the approach produced reliable whisker deflections up to 20-Hz stimulation frequency.

All stimulation protocols consisted of 16-s stimuli blocks interleaved with 44-s breaks. The stimuli analyzed here were given at either 1, 5, 9, 13, or 17 Hz frequency. Additionally, 3 Hz and 7 Hz stimuli were given under anesthesia but were not included in the current study since these frequencies were left out from awake experiments to minimize the scanning time. In a single experiment, each frequency was repeated four times, leading to 20 1-min stimulus and baseline blocks per scan for the current analysis. The order of the different frequencies and their repetitions within each experiment was randomized, as was also the order of mechanical and electrical protocols in the experiments conducted under anesthesia.

### 5.5 Isoflurane + medetomidine anesthesia protocol

In the experiments done under anesthesia, rats were lightly anesthetized with a combination of isoflurane (Attane vet 1000 mg/g, Piramal Healthcare UK Limited, Northumberland, UK) and medetomidine (Domitor vet 1 mg/ml, Orion Corporation, Espoo, Finland). At the beginning of each session, anesthesia was induced with 5% and maintained with 2 % isoflurane during the positioning of the rat into the custom-made MRI holder. A stainless-steel cannula was placed subcutaneously in the back of the animal for the delivery of medetomidine by an infusion pump (AL-1000, World Precision Instruments, Friedberg, Germany). A medetomidine bolus (0.015 mg/kg, s.c.) was given 12.9 ± 4.4 min (n = 26) after the induction of isoflurane anesthesia, followed by infusion (0.03 mg/kg/h s.c.) 15 min later. After the medetomidine bolus, isoflurane was gradually reduced and adjusted (typically 0.3-0.5 %) to achieve a respiration rate between 45 and 65 breaths per minute. The first fMRI scan started 48 ± 14 min (n = 26) after the initial medetomidine bolus, giving time for the level of anesthesia to stabilize. After finishing measurements, the rats were given atipamezole (0.5 mg/kg s.c.; Antisedan vet 5 mg/ml, Orion Corporation, Espoo, Finland) and were returned to their cages.

### 5.6 Magnetic resonance imaging

The MRI data acquisition was similar to that described earlier^44^. Briefly, data were acquired in a 9.4 T/31 cm bore magnet with 12-cm inner diameter gradient coil set, interfaced with an Agilent DirectDRIVE console (Palo Alto, CA, USA). A linear 22-mm inner diameter surface coil (Neos Biotec, Pamplona, Spain) was used for both transmission and receiving the radio-frequency signals.

Anatomical reference images were acquired with an MB-SWIFT sequence with the following parameters: 4000 spokes per spiral, 16 stacks of spirals, repetition time 3 ms for a spoke, 4 radiofrequency pulses per spoke, flip angle 5-6°, excitation/acquisition bandwidths of 192/384 kHz, matrix size 256^3^, field of view 40^3^ mm, and 156 µm isotropic resolution. To increase anatomical contrast, a magnetization transfer pulse (γB1 = 125 Hz, offset 2000 Hz, duration of 20 ms) was given at every 32 spokes. The sequence had a total acquisition time of ∼4 min.

Functional data were obtained with an MB-SWIFT sequence with the following parameters: 2047 spokes per spiral, 1 spiral, repetition time 0.97 ms for a spoke, 4 radiofrequency pulses per spoke, flip angle 5-6°, excitation/acquisition bandwidths of 192/384 kHz, matrix size 64^3^, field of view 40^3^ mm, and 625 µm isotropic resolution. The acquisition time for a single fMRI volume was 2 s. The first stimulus block was preceded by a 3-min baseline period to ensure the stabilization of baseline. Subsequently, the lengths of awake and anesthesia fMRI scans were 690 (23 min) and 930 volumes (31 min), respectively. The first two volumes were discarded as the signal was reaching a steady-state.

A physiology monitoring equipment (Model 1025, Small Animal Instruments Inc., New York, NY, USA) was used to follow the breathing rate in all animals. In anesthetized animals, the breathing rates were 48 ± 9 breaths per minute (n = 52) at the onset of the fMRI scans. In awake animals, the breathing rates were 108 ± 14 breaths per minute (n = 28) at the onset of the scan and varied up to 144 ± 22 breaths per minute during the 23-min fMRI measurement. Additionally, the temperature of anesthetized rats was monitored with a rectal probe. A warm water circulation system (Corio CD, Julabo, Seelbach, Germany) was used to keep the animals warm (36.9 ± 0.6 °C, n = 51, measured at the onset of the anesthesia fMRI scans). A video camera (12M-i, MRC Systems GmbH, Heidelberg, Germany) was used to visually monitor the whisker deflections and animal behavior during the MRI.

### 5.7 Data processing and analysis

Generally, the data processing and analysis shared similarities with our previous work^44,45^. A Snakemake^85^ (https://snakemake.readthedocs.io) script was used to perform the steps of 1) reconstructing images into NIfTI, 2) saving data in the BIDS structure^86^ (https://bids.neuroimaging.io), 3) co-registration, and 4) motion correction. The MB-SWIFT^87^ data were reconstructed using an in-house Python (version 3.10) script for a center-out data re-gridding and iterative FISTA algorithm^88^, performed volume-by-volume with 13 iterations. The reconstructed anatomical brain volumes were co-registered to a study-specific template using N4 bias correction^89^, and linear and non-linear SyN registrations of Advanced normalization tools^90^ (ANTs, http://stnava.github.io/ANTs).

Even though the rats were head-fixed, which diminished movement of the skull, the fMRI data were motion-corrected with a tailor-made Python script using masking, N4 bias correction, and ANTs rigid co-registration of each volume to the first volume of the 4-D data set. Motion correction parameters were saved for further use. The motion-corrected fMRI images were mirrored in the left-right orientation, if needed, to locate the stimulation in the same side in all data sets. Subsequently, the fMRI images were translated to the template reference frame for group-level analyses using linear and non-linear transformations from the anatomical co-registration. No spatial smoothing or global signal regression was applied to the fMRI data.

The first-level statistical maps for each dataset were computed using FLAME from FSL FEAT^91^. A mask covering the whole brain was used. The parameters used for the hemodynamic response function with the general linear model were derived in separate experiments (Valjakka et al., submitted). At the individual experiment level, each stimulation frequency was treated as an event, resulting in four stimulus blocks for each event. Motion parameters from the rigid motion correction were used as confounding regressors. The group-level maps were produced by concatenating all experiment level maps and testing if the responses to stimuli within the nine subgroups, namely low-, mid-, and high-frequency stimuli in the three experimental groups, differed statistically from zero. The statistical tests were performed with the Randomise^92^ tool while taking multiple comparisons into account by using the family-wise error (FWE) correction. FreeSurfer (Freeview, version 2.0, https://surfer.nmr.mgh.harvard.edu/) and FSLeyes (FSL version 6.0.5, https://fsl.fmrib.ox.ac.uk/fsl/fslwiki) were used for the evaluation and representation of the statistical maps.

ROIs were drawn with FSLeyes on the fMRI reference frame according to an anatomical atlas^51^, aiming to exclude any contribution from ventricles and large blood vessels. Regions were taken for further analyses if they included significant (p<0.005, FWE-corrected) voxels with any stimulation frequency range or experimental group. To minimize the potential effect of motion, the fMRI time series were regressed using motion correction parameters. Subsequently, ROI time series were computed as an intensity average under the ROI, temporally smoothed with a 0.4 smoothing parameter of smoothness priors detrending method^93^, and grouped according to different frequency ranges and experimental groups.

The average response values for each ROI were calculated from a 11-volume (22-s) window starting at the onset of the 16-s stimulus. An exception of a shorter 5-volume (10-s) window was used for SC due to its complex biphasic response shape, where the 11-volume window resulted in close-to-zero values, not reflecting the underlying response. The stimulation frequency-dependence of fMRI responses was investigated by fitting a linear function on the average responses of different stimulation frequency ranges, assuming the pools to be equidistant from each other. The slope from linear fitting was used in determining whether the fMRI response displayed an association with stimulation frequency. The linearity of association was determined by testing the normality of the residuals. Lastly, the categorization of different fMRI response profiles and temporal signal characteristics were obtained by using hierarchical clustering with the linkage-function in the Python scipy-module. Clusters consisting of a single region and high hierarchy were excluded from discussion.

Group-level values are represented as group-level mean ± standard deviation unless stated otherwise. The types of statistical tests used are mentioned in the Results section and in the figure legends. The threshold for statistical significance was set to p < 0.005 in voxel-wise statistical maps, and elsewhere to p < 0.05, unless otherwise stated. Corrections for multiple comparisons were performed with FWE-correction, unless stated otherwise.

## 6. Acknowledgements

The authors thank Irina Gureviciene, Maarit Pulkkinen, and Petteri Stenroos for help in the head implant surgeries. This work was supported by NIH grants P41 EB027061 and R01 MH127548-01, Research Council of Finland grant 355391, Sigrid Jusélius Foundation, Kuopio Biomedical Imaging Unit (part of Biocenter Kuopio, Finnish Biomedical Imaging Node, and EuroBioImaging), and Phenotyping Center (part of Biocenter Kuopio). The computation was performed on servers provided by UEF Bioinformatics Center, University of Eastern Finland, Finland

## 7. Author contributions

J.P, J.V, H.T, S.Mi, S.Ma, and O.G designed the study. J.P and J.V conducted the in vivo work. R.S, S.Mi, and S.MA assisted in conceptualization and methodology. J.P, J.V, R.S, E.P, and O.G participated in data processing, analysis, and interpretation. J.P and O.G supervised the work. J.P wrote the original draft. H.T, S.Mi, S.Ma, and O.G contributed to funding acquisition and resources. O.G administered the project. All authors contributed to manuscript review and editing process.

## 8. Competing interests

The authors have no conflicts of interest to disclose.

## Abbreviations

Aud: auditory cortex
fMRI: functional magnetic resonance imaging
H6: hemisphere of cerebellar lobule 6 i.e. simplex
H7a: anterior hemisphere of cerebellar lobule 7 i.e. crus 1 and 2
H7p: posterior hemisphere of cerebellar lobule 7 i.e. paramedian 1
IC: inferior colliculus
Iso+Med: isoflurane and medetomidine anesthesia
MB-SWIFT: Multi-Band SWeep Imaging with Fourier Transformation
MGN: medial geniculate nuclei
PHC: parahippocampal cortex
Po: posterior thalamic nuclei
PPC: posterior parietal cortex
Pr5/Sp5: principal and spinal trigeminal nuclei
RSC: retrosplenial cortex
S1bf: primary somatosensory cortex, barrel field
S1lip: primary somatosensory cortex, lip region
S2: secondary somatosensory cortex
SC: superior colliculus
TeA: temporal association cortex
V7-8: vermis of cerebellar lobules 7 and 8
Vis: visual cortex
VPM: ventral posteromedial thalamic nuclei
Zero-TE: zero echo time
ZI: zona incerta

## Supplementary Tables

**Supplementary Table 1.** Numerical details related to the animals and stimulation experiments. fMRI, functional magnetic resonance imaging.

|  | Males | Females | Total |
| --- | --- | --- | --- |
| **Number of animals** | **7** | **6** | **13** |
| **Weight across fMRI sessions** | 419 ± 49 g | 302 ± 36 g | 357 ± 72 g |
| **Age across fMRI sessions** | 18 ± 5 wk | 29 ± 6 wk | 24 ± 8 wk |
| **Number of fMRI sessions** | 26 | 28 | 54 |
| **Total number of fMRI scans** | **40** | **40** | **80** |
| Electrical stimulation under anesthesia | 14 | 12 | 26 |
| Mechanical stimulation under anesthesia | 14 | 12 | 26 |
| Mechanical stimulation in awake state | 12 | 16 | 28 |
| **Total number of stimuli blocks** | **800** | **800** | **1600** |
| Electrical stimulation under anesthesia | 280 | 240 | 520 |
| Low-frequency (1 Hz) | 56 | 48 | 104 |
| Mid-frequency (5, 9 Hz) | 112 | 96 | 208 |
| High-frequency (13, 17 Hz) | 112 | 96 | 208 |
| Mechanical stimulation under anesthesia | 280 | 240 | 520 |
| Low-frequency (1 Hz) | 56 | 48 | 104 |
| Mid-frequency (5, 9 Hz) | 112 | 96 | 208 |
| High-frequency (13, 17 Hz) | 112 | 96 | 208 |
| Mechanical stimulation in awake state | 240 | 320 | 560 |
| Low-frequency (1 Hz) | 48 | 64 | 112 |
| Mid-frequency (5, 9 Hz) | 96 | 128 | 224 |
| High-frequency (13, 17 Hz) | 96 | 128 | 224 |

**Supplementary Table 2.** The obtained uncorrected p-values for estimating the significance of the slope and the linearity between the average functional magnetic resonance imaging response and the stimulation frequency. For each region-of-interest and group, we tested whether the slope between stimulation frequency and average response deviated from 0 (t-test), and whether the relationship between the stimulation frequency and average response was linear (normality test for residuals of the fit). Iso+Med, isoflurane and medetomidine anesthesia.

| ROI | Iso-Med Electric |  | Iso-Med Mechanical |  | Awake Mechanical |  |
| --- | --- | --- | --- | --- | --- | --- |
| | Slope ≠ 0 | Non-linear | Slope ≠ 0 | Non-linear | Slope ≠ 0 | Non-linear |
| Pr5 / Sp5 | **<0.0001** | 0.5273 | **0.0004** | 0.3473 | **0.0058** | 0.3701 |
| VPM | **0.0058** | **0.0033** | **0.0024** | **0.0152** | **0.0060** | 0.1559 |
| PO | 0.5150 | **0.0402** | 0.3662 | 0.3134 | **0.0260** | 0.9527 |
| S1bf | **0.0037** | **0.0032** | **<0.0001** | **<0.0001** | **<0.0001** | 0.4061 |
| S2 | 0.0758 | **0.0007** | 0.4466 | **<0.0001** | **0.0001** | 0.0754 |
| S1lip | **0.0007** | **<0.0001** | **0.0031** | **<0.0001** | **0.0004** | 0.3990 |
| Aud | **0.0028** | **0.0476** | 0.0648 | **0.0015** | 0.0567 | 0.0784 |
| Vis | 0.4019 | **0.0023** | 0.6921 | **<0.0001** | **0.0341** | 0.3534 |
| IC | 0.7517 | 0.3994 | **0.0135** | **0.0420** | **0.0005** | 0.3138 |
| SC | 0.0692 | 0.2036 | 0.9277 | 0.1569 | 0.7407 | 0.7963 |
| MGN | 0.9052 | **0.0076** | 0.0956 | **0.0229** | 0.6573 | 0.8440 |
| ZI | **0.0024** | 0.0522 | 0.4527 | 0.3005 | 0.9344 | 0.9834 |
| RSC | 0.5449 | 0.6386 | 0.9021 | 0.5099 | 0.1413 | **0.0066** |
| TeA | 0.9842 | 0.5479 | 0.8334 | 0.1029 | 0.2992 | 0.9300 |
| PPC | 0.4112 | **0.0023** | 0.7417 | **0.0186** | **0.0333** | 0.1055 |
| PHC | 0.7726 | **0.0009** | 0.6023 | 0.1251 | 0.6857 | 0.2608 |
| H6 | **0.0458** | 0.8417 | 0.0501 | 0.1940 | 0.7246 | **0.0253** |
| H7a | 0.060 | 0.0925 | 0.3478 | **0.0307** | **0.0096** | 0.0644 |
| H7p | 0.3823 | 0.4145 | 0.7664 | 0.2291 | **0.0425** | 0.4426 |
| V7-8 | 0.1350 | 0.2299 | 0.8871 | 0.3868 | **0.0277** | 0.3434 |

## Supplementary Figures

**Supplementary Figure 1.**
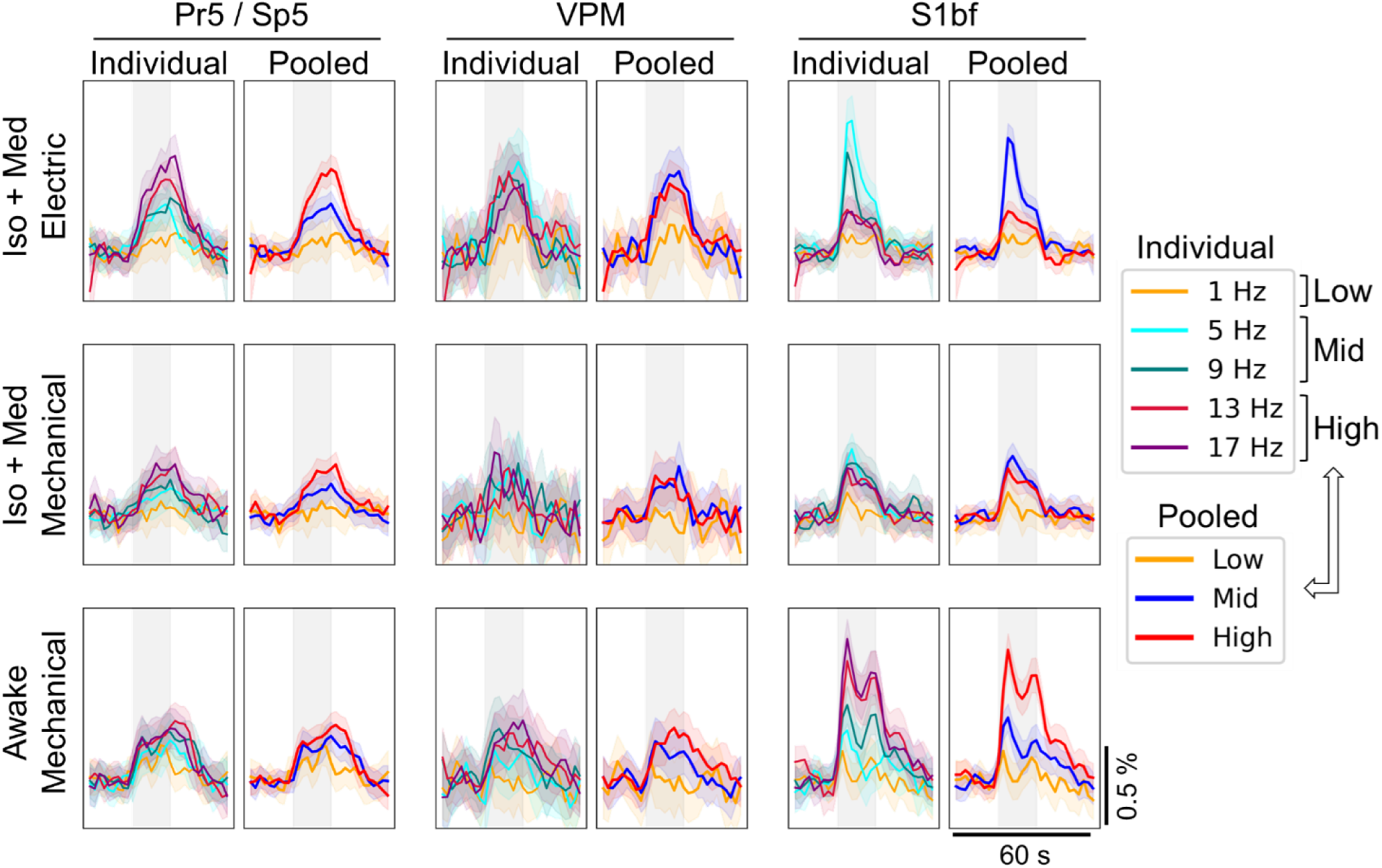
Frequency-specific and pooled time series obtained from the key nodes of the whisker-mediated tactile system. Because of the similarity of the responses to certain stimulation frequencies, their correspondence to natural whisking frequencies, and to simplify the representation of complex and multilayered data, the five different stimulation frequencies were pooled into three subgroups. Each frequency-specific and pooled mean time serie consists of 104-112 or 208-224 repetitions, respectively. The shaded vertical gray region indicates the timing for the 16-s stimulus block. The 90% confidence interval is shown as a shaded region around the mean time series. Pr5/Sp5, principal and spinal trigeminal nuclei; S1bf, primary somatosensory cortex barrel field region; VPM, ventral posteromedial thalamic nuclei.

**Supplementary Figure 2.**
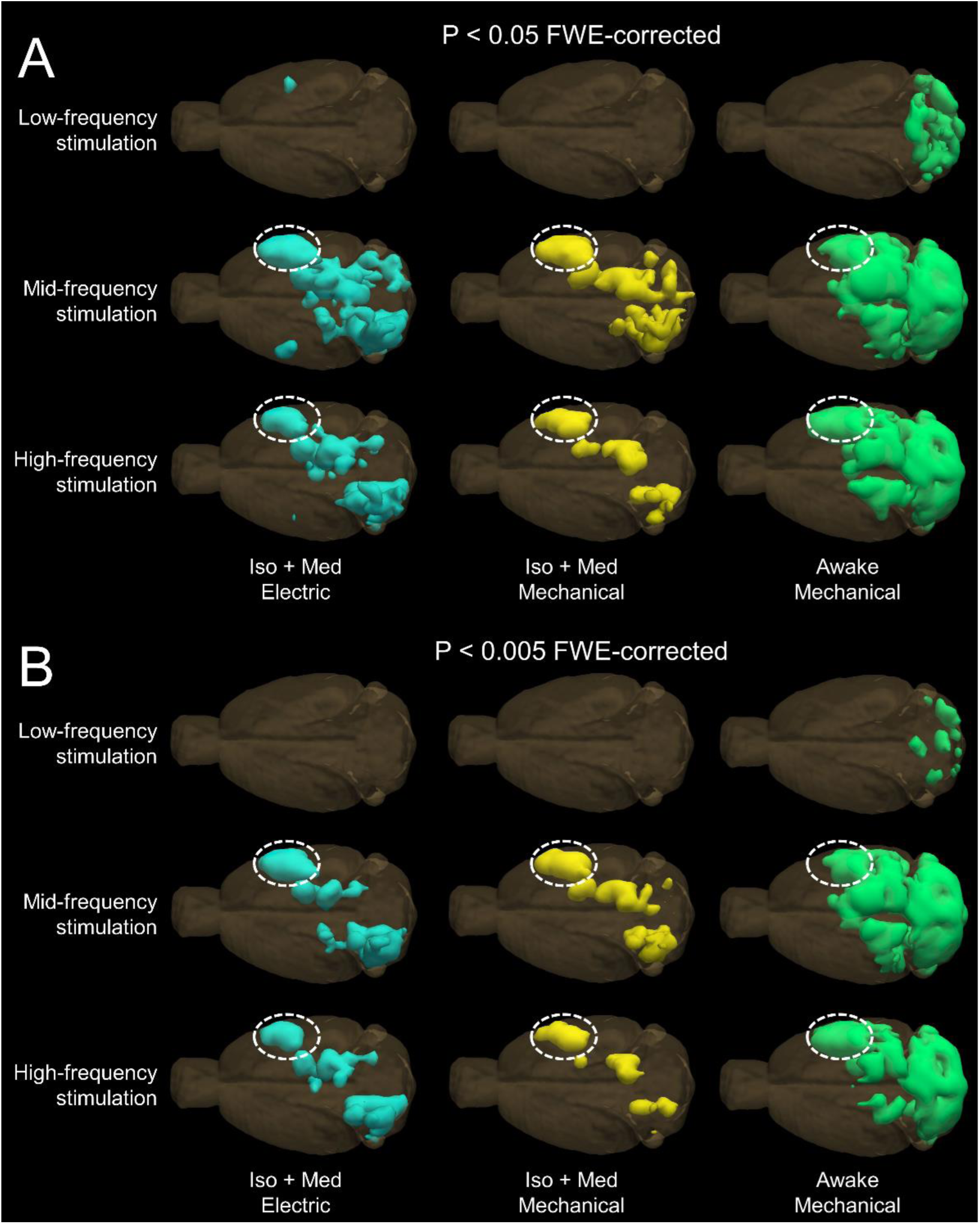
3-D illustration of significant voxels (A, p<0.05, FWE-corrected; B, p<0.005, FWE-corrected) obtained with low-, mid-, and high-frequency stimuli in each group. The results are obtained from 104-224 stimuli blocks per map. Generally, low-frequency stimuli produced the spatially smallest responses, and the localization of signal changes to mid- and high-frequency stimuli resembled each other. However, there were also clear differences in the statistical maps to the mid- and high-frequency stimuli. As an example, the high-frequency stimulus yielded spatially more widespread signal changes within the S1BF (dashed ellipsoid) in awake rats [83 vs 63 significant voxels (p<0.005, FWE-corrected)], but more restricted signal changes in anesthetized rats with both electrical [43 vs 67 significant voxels (p<0.005, FWE-corrected)] and mechanical [50 vs 70 significant voxels (p<0.005, FWE-corrected)] stimuli. These findings suggest that anesthesia can affect the relationship between stimulation frequency and the spatial extent of the fMRI responses within regions. Iso+Med, isoflurane and medetomidine anesthesia.

## Notes

### Competing Interest Statement

The authors have declared no competing interest.

